# Reconciling fish and farms: Methods for managing California rice fields as salmon habitat

**DOI:** 10.1101/2020.08.03.234062

**Authors:** Eric J. Holmes, Parsa Saffarinia, Andrew L. Rypel, Miranda N. Bell-Tilcock, Jacob V. Katz, Carson A. Jeffres

## Abstract

The rearing habitat for juvenile Chinook Salmon (Oncorhynchus tshawytscha) in California, the southernmost portion of their range, has drastically declined throughout the past century. Recently, through cooperative agreements with diverse stakeholders, winter-flooded agricultural rice fields in California’s Central Valley have emerged as promising habitat for rearing juvenile Chinook Salmon. From 2013 to 2016, we conducted a series of experiments examining methods for rearing fall-run Chinook Salmon on winter-flooded rice fields in the Yolo Bypass, a modified floodplain of the Sacramento River in California. These included: 1) influence of field substrate differences from previous season rice harvest; 2) effects of depth refugia from avian predators (trenches); 3) field drainage methods to promote efficient egress of fish; and 4) in-field salmon survivorship over time. Zooplankton (fish food) in the winter-flooded rice fields were 53-150x more abundant when directly compared to the adjacent Sacramento River. Correspondingly, somatic growth rates of juvenile hatchery-sourced fall-run Chinook Salmon stocked in rice fields were two to five times greater versus fish in the adjacent Sacramento River. Post-harvest field substrate treatments had little effect on the lower trophic food web and had an insignificant effect on growth rates of in-field salmon. Though depth refugia did not directly increase survival, it buffered maximum water temperatures in the trenches and facilitated outmigration from fields during draining. Rapid field drainage methods yielded the highest survival and were preferable to drawn-out drainage methods. High initial mortality immediately after stocking was observed in the survival over time experiment with stable and high survival after the first week. In-field survival ranged 7.4–61.6% and increased over the course of the experiments. Despite coinciding with the most extreme drought in California’s recorded history, which elevated water temperatures and reduced the regional extent of adjacent flooded habitats which concentrated avian predators, the adaptive research framework enabled incremental improvements in design to increase survival. The abundance of food resources and exceptionally high growth rates observed during these experiments illustrate the benefits associated with reconciling off-season agricultural land use with fish conservation practices. Without any detriment to flood control or agricultural yield, there is great promise for reconciliation ecology between agricultural floodplains and endangered fish conservation where minor alterations to farm management practices could greatly enhance the effectiveness of fish conservation outcomes.

## Introduction

Reconciliation ecology, or the modification of human dominated ecosystems for wildlife conservation is becoming increasingly important, especially in freshwater ecosystems [1,2]. Indeed, streams, rivers and their associated floodplains are some of the most altered ecosystems in the world, and reconciliation ecology will be increasingly needed to preserve endangered species in degraded environments, and assist in recovering declining fauna [3–5]. Given human population expansion and the concurrent growth of agriculture, multi-benefit land use agreements are needed that align with the needs of societies as well as ecosystems [6]. Agricultural floodplains are an ideal location for case-studies on innovative arrangements since they are managed to perform economically valuable functions of human food production and flood risk mitigation as well as still providing passive benefits through the breakdown of organic material, nutrient cycling, aquifer recharge, habitat creation, and conservation of biodiversity in heavily altered landscapes [7,8]. Though river floodplain environments are largely in decline globally [9,10], floodplain-oriented reconciliation ecology is especially attractive because exposure of large numbers of fish to abundant food resources and habitats similar to those in which native species are adapted may benefit survival and fitness [11,12]. The culturing of fish in rice fields has successfully taken place for thousands of years in East Asia during both the growing season and off-season, providing a valuable protein resource, natural fertilizer for agricultural fields, and refugia/food for native fishes [13,14]. In North America, much of the existing reconciliation work in agricultural floodplains has focused on waterfowl conservation [15,16]; thus applications for fishes have lagged behind. Resultantly, there are myriad important but fine-detailed questions regarding how to best prepare agricultural floodplains to enable fish conservation.

*Oncorhynchus tshawytscha*, commonly known as Chinook Salmon, are declining in California [17]. A conservative pre-European establishment population estimate in the Central Valley fish was 2 million annual adult spawners, and sustained a sizable commercial ocean fishery [18]. Anthropogenic causes of its decline are related to a combination of overfishing, dam and levee construction, water diversions, land use changes, logging, hatcheries, and climate change [19,20]. Current restoration efforts designed to improve spawning conditions and produce more juvenile salmon include managing for minimum instream flows, cold water releases, gravel enhancement, trap-and-haul programs, and hatchery production [21,22]. However, the ultimate success of these efforts, measured by resulting adult returns, depends largely on freshwater rearing conditions and the correlated early ocean success of juveniles [23].

California’s Central Valley floodplain is an optimal region for practicing floodplain reconciliation ecology due to the amount of wetland habitat that has been lost in this region through the past century. Prior to the mid-1800s, there were an estimated 4 million acres of seasonal wetlands in California’s Central Valley [24], which provided floodplain habitat with an abundance of food resources juvenile Chinook Salmon. Of the historic wetland habitats in California, approximately 95% of floodplain habitat has been disconnected from rivers by levees and channelization, drastically reducing quality rearing conditions for out-migrating salmon [25,26]. Much of the remaining wetland habitat accessible to juvenile salmon is highly altered, confined to flood bypasses, and designed to drain flood waters rapidly to protect cities while accommodating agricultural production in the dry season. Though much of the historical alluvial floodplain in California is inaccessible to salmon with remaining areas confined to primarily agricultural lands in the Yolo Bypass, Sutter Bypass, and other parcels located within Army Corps river levee system, many properties of productive seasonal wetlands persist, presenting opportunities for conservation. In particular, winter-flooded rice fields develop high densities of invertebrate prey that promote fast juvenile Chinook Salmon growth rates in addition to environmental conditions that resemble natural habitat conditions [27–30].

Using irrigation infrastructure to flood large areas within existing migratory routes during the winter non-growing season presents a potential pathway to expanding the extent and enhancing the quality of rearing habitat available to the salmon populations of the Central Valley. Managed agricultural floodplains have the potential to supply juvenile fish with high quality floodplain habitat and abundant natural food sources before downstream migration [29,31]. These benefits accrue with prolonged duration of floodplain inundation with extended flooding during the winter and early spring seasons allowing for the development of invertebrate prey and improved foraging opportunities for fish [32]. Vigorous somatic growth of juvenile salmon is typically observed in fish rearing under these floodplain conditions, leading to potential survivorship benefits during outmigration [30,33]and their critical transition to the ocean environment [34,35]. While the potential benefits to juvenile Chinook Salmon rearing in flooded rice fields is well established, there is little research testing methodologies for establishing the optimal physical and biological conditions to achieve maximal benefit in managed agricultural floodplains.

The primary goal of this study was to compare potential farm management practices intended to improve habitat conditions and growth and survival of juvenile Chinook Salmon within winter-flooded post-harvest rice fields on the floodplain of the Central Valley of California. Data on which methods provide the best conditions for juvenile salmon could be useful to growers interested in participating in fish conservation activities and resource managers developing guidance for multi-benefit land use intended to improve the habitat quality of managed floodplains for salmon and other native fishes of conservation concern. Starting in 2013, we utilized an adaptive research framework to test rearing methodology of juvenile salmon yearly until 2016. In 2013, our objective was to investigate food resources (invertebrate prey) and salmon growth response to rice field substrate types (comparing post-harvest field preparation methods). In 2014, we investigated the role that depth refugia played in determining in-field survival and outmigration behavior. In 2015, water drainage practices were investigated to determine if natural floodplain hydraulics could be mimicked to maximize efficient egress of juveniles upon field draining. In 2016, we investigated in-field survival over time to evaluate optimal rearing durations.

## Methods

### Study area

Experiments took place in the Yolo Bypass, a 24,000-ha flood bypass along the Sacramento River in California, USA. Nine 0.81 ha replicated fields were constructed on Knaggs Ranch—a farm predominantly producing rice (Fig 1). An inlet canal routing water from the Knights Landing Ridgecut Canal independently fed each of the nine fields, and all fields drained into an outlet canal. The outlet canal ultimately emptied into the Tule Canal, which runs north to south along the east side of the bypass. Each field had rice boxes (structure using stacked boards to control water elevation and flow) on the inlet and outlet of each field. Water depths, as measured in the middle of the fields, were maintained between 0.3m to 0.5m for all years. Inlet structures were fitted with 4mm mesh screens to permit water inflow and prevent egress of stocked salmon. Outlet structures were fitted with mesh screens in the 2013 and 2016 experiments. However, in 2014 and 2015, outlet structures were left open with a 5-cm diameter hole drilled in the middle of a 3.8cm × 14cm board and placed near the top of the water level in the rice box to investigate volitional outmigration patterns of the stocked salmon. Each outlet structure was fitted with a live car trap placed in the outlet canal, which allowed for collection of all exiting fish. In 2014 and 2015, live cars were checked daily for the duration of the experiments to enumerate the number of emigrating salmon. In past experiments we observed a tendency for a portion of hatchery fish to “scatter” upon initial release into floodplain fields. This behavior reliably abated after several days as fish acclimated to new conditions. For this reason, downstream exiting fish were restocked back to the inlet side of the fields for the first week of 2014. In 2015, fish were similarly restocked for 2 weeks.

**Fig 1.**
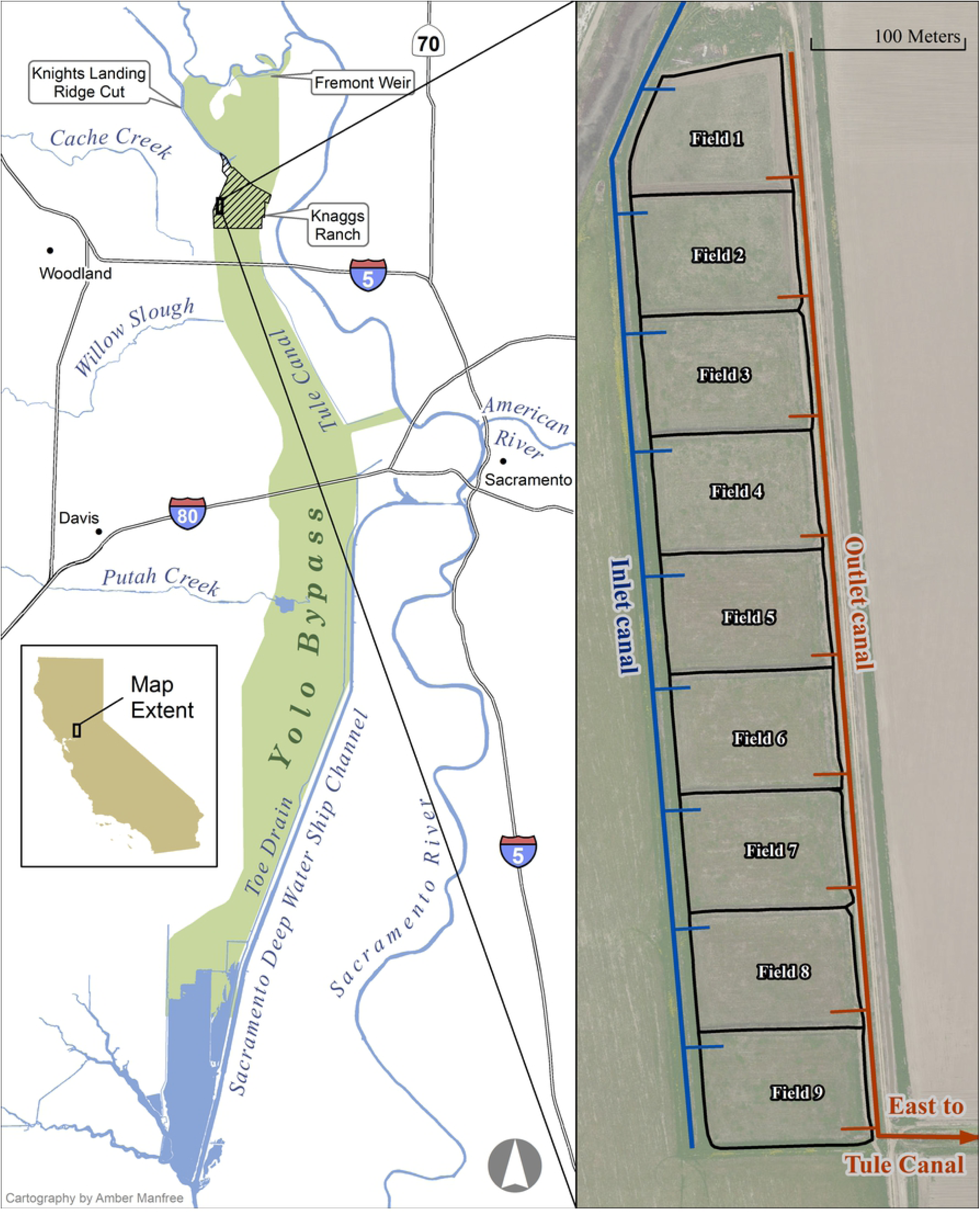
Map of the study area. Geographic extent of the Yolo Bypass of California (left) with a detailed view of the nine experimental rice fields (right).

### Experiments

#### Substrate type – 2013

After harvest, rice farmers remove residual rice straw remaining in the fields using one of several methods; thus an important question was whether differences in treatment of rice straw created different outcomes for rearing fish. In 2013, nine fields were randomly assigned to one of three post-harvest substrate treatments: rice stubble, disced, or fallow. The rice stubble substrate treatment consisted of standing stalks (heights ranging from 0.23-0.35m) that remained after rice plants were cut for harvest using a rice harvesting combine tractor. The disced treatment consisted of plowing rice straw into the soil, a practice farmers use to promote stubble decomposition. The fallow habitat had not been planted with rice during the previous growing season but instead consisted of weedy herbaceous vegetation that voluntarily colonized the fields during the growing season and was left standing during the experiment.

The 2013 experiment was described in detail by our colleagues [27] from the lower trophic food web perspective and referenced in a forthcoming overview of research on salmon in farm fields [36]. The methods are briefly summarized above to provide context for subsequent years of the evolving project which was built on an adaptive research framework.

#### Depth refugia – 2014

Avian predation on fish in aquaculture fields is a well-known problem [37–39]. Avian predation has potential to be a significant source of mortality on fish in winter-flooded rice fields as California’s Central Valley is positioned directly within the winter habitat of diverse bird populations in the Pacific flyway [40,41]. We evaluated trenching as one method for reducing potential avian predation on fish in winter-flooded rice fields. In 2014, nine fields all with a disced substrate, were randomly assigned to one of three treatments: three fields were assigned no perimeter trench, three were assigned a 0.5m deep perimeter trench, and three were assigned a 1.0m deep perimeter trench. All trenches were constructed on the north and east sides of the fields running continuously from the inlet structure in the northwest corner to the drain structure in the southeast corner. All trenches were approximately 1.0m wide with the outermost edges of the trench spaced approximately 1.0m from the exterior levee surrounding the field. We created this spacing specifically so depth refuges were outside the striking distance of wading birds such as herons and egrets that frequent the shallow water of the perimeter levees. Survival data for 3 fields was excluded from the analysis due to loss of containment on the inlet side of three fields (fields 3, 4, and 7, one from each treatment) during the last week of the experiment allowing fish to escape upstream into the inlet canal. Ancillary effects of the trench treatments including field drainage efficiency and volitional migration patterns were investigated between treatments.

#### Drainage practices – 2015

Floodplain hydrology provides important cues for movement and egress of floodplain species [42,43]. Managing winter-flooded rice field habitat for fishes may require the manipulation of draining hydrology to maximize survival and volitional egress of fish off fields. These dynamics could be especially important in rice fields where intermittent floods could entrain wild salmon and other species within fields (e.g., in the Yolo or Sutter Bypasses of the California Central Valley). To investigate drainage practice effects on fish survival, the nine fields were randomly assigned one of three draining treatments: 1) fast drain, where inlet water was cut off and outlet boards were removed rapidly, resulting in the water draining off the fields in a single day; 2) slow drain with inflow, where water levels were lowered by 5 cm per day at the outlet while inflow was maintained through a mesh screen; and 3) slow drain without inflow, where water levels were lowered 5 cm per day at the outlet and inflow was cut off by boarding up the inlet structure. The drainage duration for slow drain procedures lasted for 10 days with daily outmigration of salmon measured in the outlet traps. All nine experimental fields had a rice stubble substrate following the rice harvest in fall 2014, and a 0.5m deep perimeter trench was constructed in all fields connecting the inlet and outlet structures running along the north and east sides of the fields. The trenches were approximately 1.0m wide and spaced 1.0m infield from perimeter levees.

#### Survival over time – 2016

Because spatiotemporal variation in predation risk on fish and thermal-oxygen conditions can be significant (especially in late winter and early spring), in-field survivorship of fish remains a concern. During 2016, to examine in-field survivorship of juvenile salmon over time, fish were stocked in six of the nine flooded experimental fields. During each of six weeks following stocking, one randomly selected field was drained until the end of the 6-week experiment. The fast drain procedure, as detailed in the 2015 experiment, was used in all fields. All fields had fallow habitat due to lack of water allocation for the 2015 growing season during an ongoing drought, and all fields had 0.5m deep trenches carried over from the previous 2015 experiment. An impending bypass flood event near the end of the study forced the drainage of the last field 4 days earlier than scheduled.

#### In-field water temperature

Water temperature in shallow inundated fields could be a stressor to juvenile salmon during warm weather periods with potential effects of slower growth, reduced smolting indices and increased predation vulnerability [44]. Across all years and fields, we recorded continuous water temperatures in 15-min intervals using HOBO U22 temperature loggers (Onset Computer Corporation, Bourne, Massachusetts, USA) anchored in a fixed vertical position on a metal t-post approximately 10cm above the substrate in the middle of each field as well as trench substrate for a representative set of treatments when applicable.

#### Zooplankton abundance

Throughout all years, a randomly stratified subset of three fields was sampled for zooplankton weekly except in 2013 where all nine field were sampled weekly. A 30-cm diameter 150-µm mesh zooplankton net (with the exception of the 2016 experiment, which used a 15-cm diameter 150-µm mesh net) was thrown 5 m and towed through the water column four times, once in each cardinal direction. In 2013, benthic macroinvertebrates were sampled separately using benthic sweeps, but due to high sedimentation, high spatial and temporal sample replication, and low overall contribution to the invertebrate community, the additional processing was deemed unnecessary in subsequent years. Furthermore, the zooplankton tow method is effective for assessing pelagic zooplankton and macroinvertebrate community assemblages while improving sample processing efficiency since it avoids the heavy sedimentation associated with benthic sweeps on wetland substrates [45]. Additionally, we also relied on the stomach contents of in-field salmon to better inform the assemblage of macroinvertebrates present in the floodplain food web and their contribution to the diet of in-field salmon (methods in next section). Sampling locations were determined randomly within the test plots via a selection of random x and y distances from a random number table. All samples were preserved in a solution of 95% ethanol. Organisms were identified with the aid of a dissecting microscope at four times magnification to the lowest taxonomic level possible using several widely recognized keys [46– 48]. Abundance estimates were calculated from homogenized subsamples of known volume and extrapolated to the volume sampled during the initial net throws.

#### Salmon stomach contents

Stomach contents of sacrificed salmon captured during seining (2013–2015) and sequential field draining (2016) were dissected for diet analysis. A total of 532 salmon stomachs (2013: n = 268, 2014: n = 144, 2015: n = 90, 2016: n = 30) were dissected using a dissecting microscope at four times magnification. Prey items were enumerated, but due to partial decomposition, prey item identification in the stomachs was limited to taxonomic order.

#### Overall salmon survival and growth

Estimates of initially stocked salmon in each field were calculated by establishing a fish per kilogram ratio and multiplying by the total weight applied to each field, except in 2016 where the overall number of stocked fish was sufficiently low to count individually (Table 1). Fish lethally sampled for stomach content analysis during weekly sampling were subtracted from the initial stocking estimate. Total salmon survival in each field was cumulatively enumerated in the outlet live car traps except during the restocking phase of 2014 and 2015 when volitionally emigrating fish were restocked to the inlet side of the fields. During field drainage, seines were used to collect stranded fish out of standing water and these fish were added to the cumulative survival count from the outlet live cars with the recovery method recorded. Survival in 2015 was calculated from only the fast drain treatment fields since the drawn out drainage methods were not comparable to drainage methods in other years.

**Table 1.**
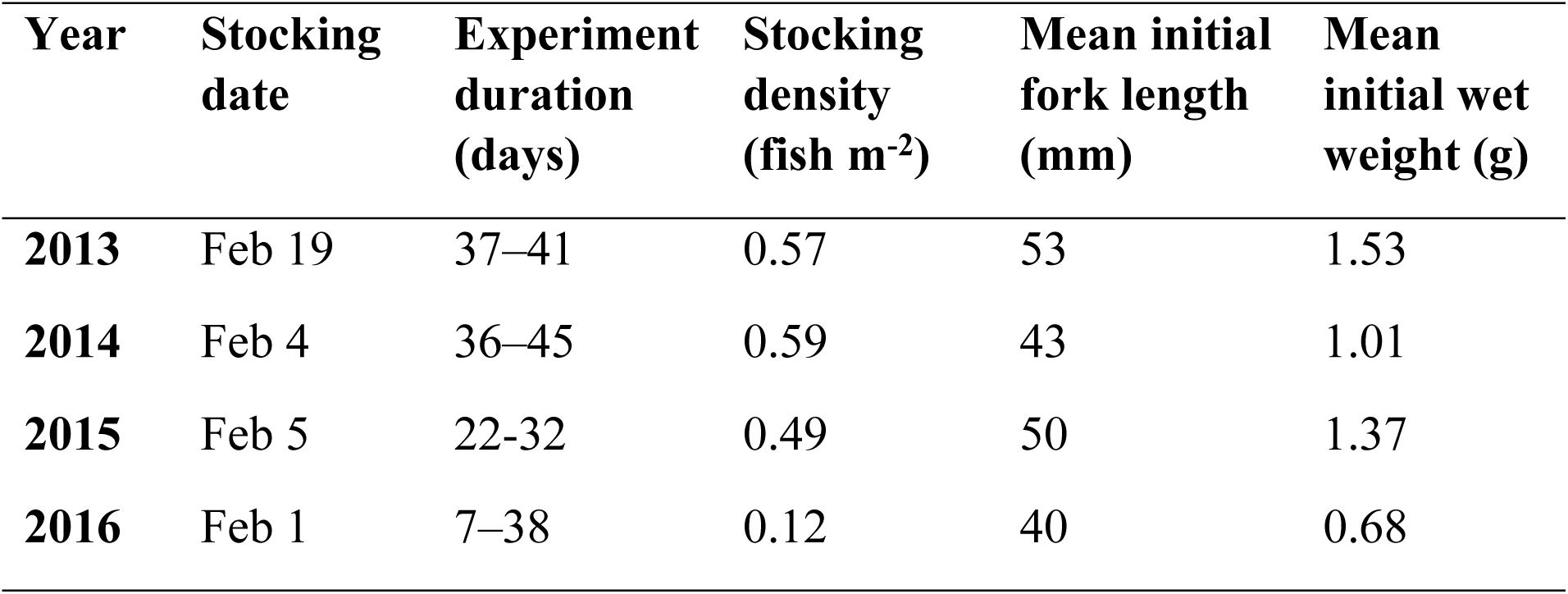
Summary of salmon stocking dates, experiment durations, stocking densities (fish m^-2^), mean initial fork length (mm), and mean initial wet weight (g).

Prior to stocking in each year, mean initial fork length and wet weight were calculated from a random sample of 30 live fish measured to the nearest millimeter and weighed to the nearest hundredth of a gram with an Ohaus Scout Pro SP202 scale (Table 1). For 2013–2015, we conducted weekly in-field fish sampling with a seine to capture a target of 30 fish per treatment, with the fork length and wet weight measured. In 2016, fish size data were collected from a random sample of 30 fish in out-migrant traps as individual fields were drained weekly.

### Statistical analysis

Percent survival for each field was calculated as the total recovered fish divided by initial stocked fish times 100. Analysis of covariance (ANCOVA) was used to test for interaction effects between field substrate treatment and time which would indicate treatment effects on salmon growth rates. In this model, fork length was the dependent variable with field substrate, day of the experiment and an interaction term as the independent variables. When the assumptions of normality and homogeneity of variance were satisfied, as tested by the Shapiro-Wilk and Levene tests respectively, a one-way analysis of variance (ANOVA) was used to test for significant differences in survival due to field drainage treatments. A post hoc Tukey honestly significant differences (HSD) test was used to test all pairwise comparisons of field drainage practices. When the assumptions of normality and/or homogeneity of variance were not satisfied, non-parametric Kruskal-Wallis analysis was used to test for significant differences in survival and daily volitional outmigration due to field trench depth treatments. A post hoc Dunn’s test was used to test all pairwise comparisons of daily volitional outmigration due to field trench depth treatments. Linear regression was used to estimate apparent growth rates and to examine the relationship between salmon survival (dependent variable) and day of the experiment (independent variable). Linear regression was also used to evaluate the relationship between daily maximum water temperature differences in the trenches (dependent variable) and daily maximum water temperature in the middle of the field (independent variable). Statistical significance was declared at α < 0.05 level. All analyses were conducted in R v3.6.1 (R Core Team, 2019) [49].

## Results

### Substrate type – 2013

Apparent fork length growth rate for juvenile salmon did not differ significantly between treatments (ANCOVA, F = 2.16, df = 2, P = 0.11). The slopes from individual linear regressions of fork length predicted by day for each treatment resulted in estimated apparent growth rates of 1.01 mm d-1 for the stubble treatment, 0.99 mm d-1 for the disced treatment, and 0.95 mm d-1 for the fallow treatment.

Our colleagues [27] found no statistical difference between total abundance of zooplankton between treatments, but did find high overall abundance and a trend of increasing zooplankton over experiment duration. Additionally, our colleagues [27] found that across all samples, cladocera were the most abundant group of zooplankton, making up over 50% of the total zooplankton assemblage.

Cladoceran zooplankton was the most common prey item found in juvenile salmon stomach contents as this taxon comprised on average 94.0% ± 1.0% SE of the diet composition across all treatments. Chironomid midges (diptera) were the second most common prey item and comprised an average of 4.8% ± 1.0% SE of the diets. Diet composition was slightly more diverse in the fallow treatment with an average of 87.3% ± 2.6% SE percent of prey items composed of cladocerans compared with an average of 97.4% ± 1.2% SE in the disced treatment and 97.3% ± 1.0% SE in the stubble treatments. A chironomid midge hatch isolated in the southernmost field (field 9) was responsible for the increased prey diversity resulting in diets composed of an average of 69% cladocera and 30% diptera. The other two fallow replicates had an average diet composition of 96% cladocera.

### Depth refugia – 2014

Depth treatments did not have a significant effect on survival (Kruskal-Wallis, χ^2^ = 0.86, df = 2, P = 0.65). Depth treatments had a significant effect on daily volitional emigration of fish before draining (Kruskal-Wallis, χ^2^ = 14.70, df = 2, P < 0.001). A post-hoc Dunn’s test revealed that the two trenched treatments had significantly more daily volitional outmigration compared to the no trench treatment (Dunn test, 0.5m trench – no trench: P < 0.001, 1m trench – no trench: P = 0.003), but that the two trench treatments were not significantly different (0.5m trench – 1m trench: P = 0.64). The average cumulative volitional outmigration before field drainage in the two trenched treatments was 15.4% ± 5.3% SE compared to the trenchless treatment, which had 3.3% ± 1.2% SE, indicating the trenches may have functioned as a migratory pathway aiding in volitional outmigration prior to field drainage. A relatively high rate of initial volitional emigration was seen in the first week (1.5%) across all fields, followed by a much lower rate of emigration in the second week (0.2%), and steadily increasing emigration in weeks three through five (0.5%, 1.6%, 5.6% respectively). Manual fish recovery with a seine at the end of field drainage in the trenchless fields ranged between 5 to 20% of the total surviving fish compared to less than 0.5% of survivors from the trenched fields which indicated a more efficient drainage procedure in trenched fields. These findings suggested functional equivalence between the 0.5m and 1.0m trench treatments.

### Drainage practices – 2015

Average salmon survival in each of the three treatments, fast drain, slow drain with flow, and slow drain without inflow, was 43.5% ± 6.5% SE, 22.8% ± 3.0% SE, and 11.4% ± 3.1% SE respectively (Fig 2). The differences between field drainage treatments were significant (ANOVA, F = 13.15, df = 2, P = 0.01). A post hoc Tukey HSD analysis revealed that pairwise comparisons of survival in the fast drain treatment were significantly higher than either slow drain treatment (P = 0.04 and P = 0.01 for slow with flow and slow without flow treatments respectively), but differences in survival between the two slow drain treatments were not significantly different from each other (P = 0.25). Volitional outmigration patterns were similar to 2014 with relatively high initial emigration in the first week (2.7%), low emigration in the second week (0.2%), and high emigration in week three (3.8%).

**Fig 2.**
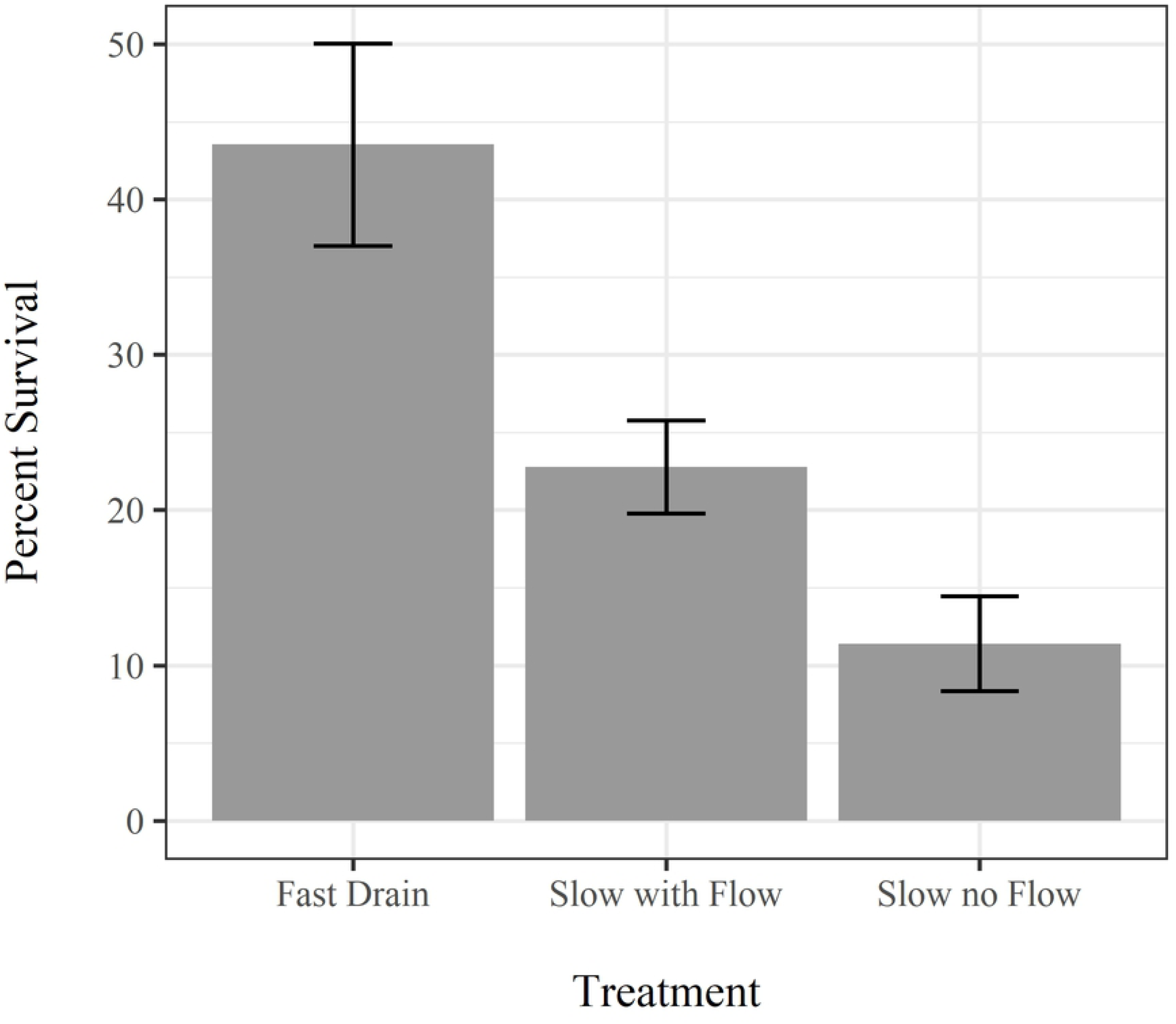
Survival response to drainage treatment. Bars represent mean percent survival between drainage treatments from the 2015 experiment. The error bars denote standard errors.

### Survival over time – 2016

Across all draining durations, in-field survival of juvenile salmon averaged 61.6% ± 6.5 SD, with the final field survival being 8.0% lower than the first field drained 31 days earlier (Fig 3). The slope of a linear survival regression model predicted by day (range: 7–38 days) was - 0.24% per day with an intercept of 67.4%. Due to low sample size (n = 6) and inherent variability in overall survival, the linear survival model had a low adjusted R^2^ and non-significant P-value (Regression, F = 1.05, df = 4, P = 0.36, R^2^ = 0.01), however, the model coefficients indicate a relatively low attrition rate (slope) after a substantial initial loss (intercept) of approximately one third during the first week of the experiment.

**Fig 3.**
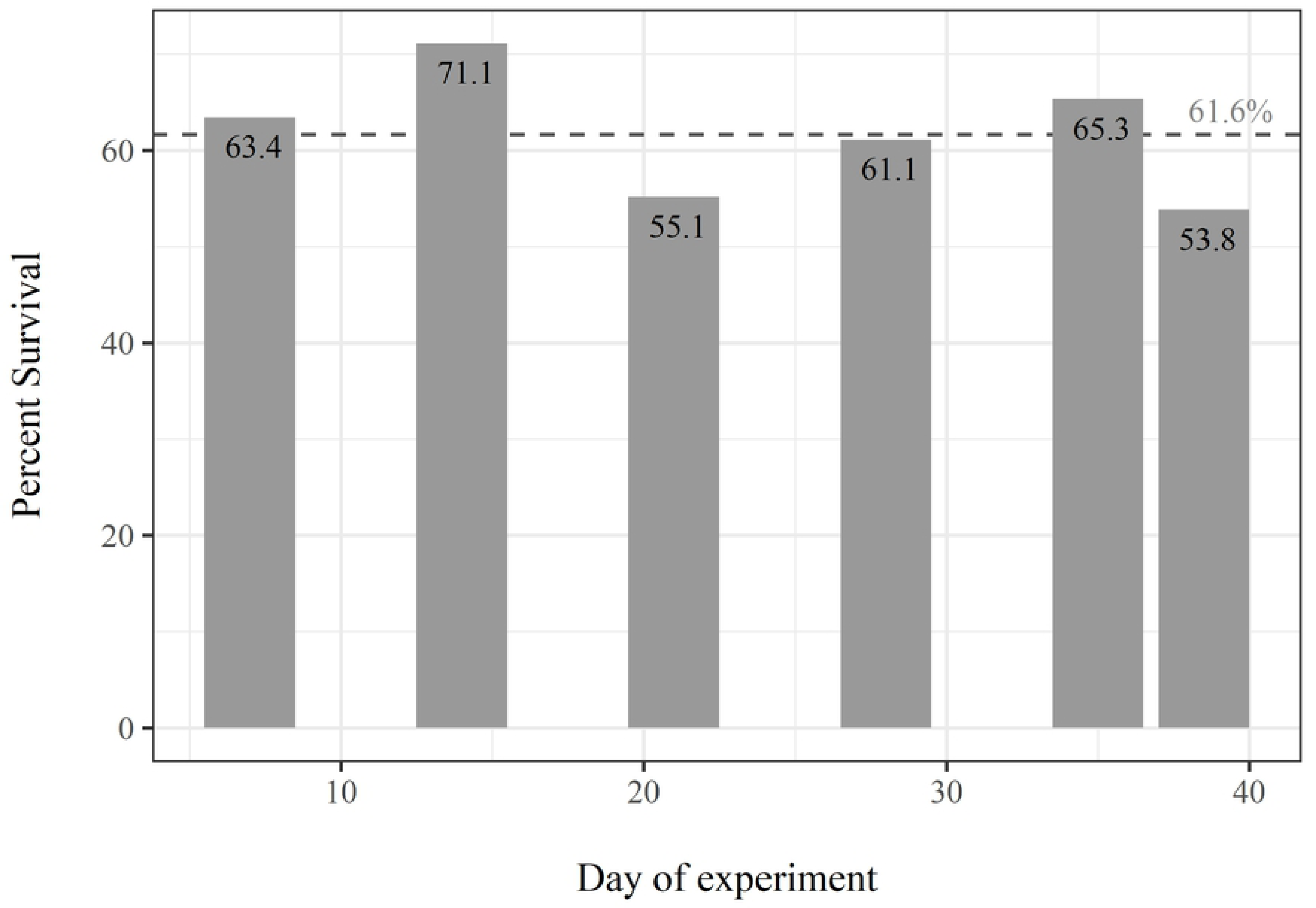
Survival over time. Bars represent percent survival of stocked salmon over time in sequentially drained fields from the 2016 experiment. The dotted line represents the mean percent survival from all fields.

### In-field water temperatures

Continuously logged water temperatures at the center of the fields during experiments from all years ranged between 5.5°C and 23.5°C. Water temperatures exceeding 21°C, which can negatively affect growth potential and predator avoidance [44], were experienced 1.6% of the time. Trenches generated thermal refugia in the bottoms of the trenches with lower daily maximum water temperatures compared to the middle of the fields by an average of 1.0 ± 0.22 SE °C and 1.8 ± 0.25 SE °C for the 0.5m and 1.0m trenches, respectively. There was a significant correlation between maximum daily water temperature in the middle of the fields and the difference between the maximum daily water temperature in the middle of the field and bottom of the 0.5m and 1.0m deep trenches (0.5m trench: regression, F = 10.4, df = 1,23, P = 0.004; 1.0m trench: regression, F = 43.89, df = 1,31, P < 0.001). Over the observed range of daily maximum water temperatures in 2014 and 2015 of 11.4 to 21.1 °C, the regression slopes showed 0.26 °C and 0.60 °C temperature reductions in the trenches per degree increase in the middle of the field for the 0.5m and the 1.0m trenches respectively.

### Overall zooplankton

Measured zooplankton densities during the experiment ranged from a low of 14,961 organisms m^-3^ on Feb 1st, 2016 in field 2 (fallow substrate) to a high of 231,966 organisms m^-3^ on Mar 20th, 2013 in field 6 (disced substrate). Overall mean annual zooplankton density was consistent each year and ranged from a low of 75,045 in 2016 to a high of 107,039 in 2015. Overall mean zooplankton densities between substrates across all years were 82,191 ± 8,697 SE, 81,283 ± 5,804 SE, and 93,585 ± 9,703 SE organisms m^-3^ for the disced, fallow, and stubble substrates, respectively. When directly compared to the adjacent Sacramento River channel habitat the managed agricultural fields had ≥ 150x zooplankton abundance in 2013 [27] and approximately 53x zooplankton abundance in 2016 [50].

### Overall salmon stomach contents

Salmon consistently showed a preference for cladoceran zooplankton as this taxon comprised >90% of the stomach contents in all years compared with ambient, in-field cladocera percent composition ranging 16.4-56.0% (Table 2, Fig 4). Mean prey organism abundance in the stomach contents for each year ranged from 158.3 ± 36.0 SE in 2016 to 278.9 ± 27.6 SE in 2013 indicating that invertebrate food resources were abundant in all years (Fig 4).

**Table 2.** Mean in-field zooplankton density m^-3^, percent cladocera from in-field zooplankton samples, and percent cladocera in salmon diets.

| Year | Mean total zooplankton density (m<sup>-3</sup>) | Mean percent cladocera in fields | Mean percent cladocera in diets (%) |
| --- | --- | --- | --- |
| 2013 | 81,417 | 56.0% | 94.0% |
| 2014 | 96,768 | 52.9% | 95.0% |
| 2015 | 107,039 | 25.2% | 92.5% |
| 2016 | 75,045 | 16.4% | 91.2% |

**Fig 4.**
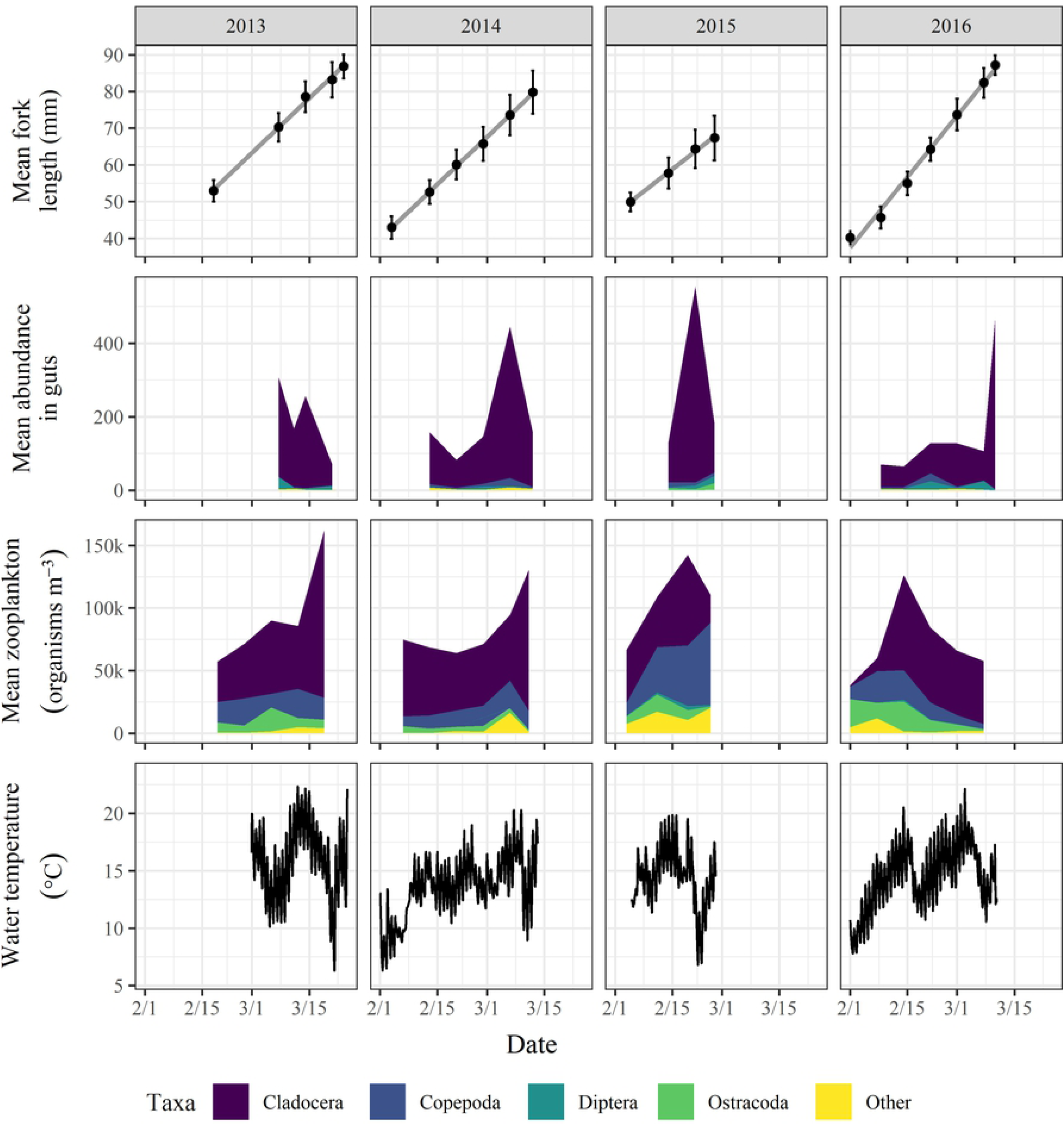
Salmon growth, stomach contents, zooplankton resources, and water temperatures in winter-flooded rice fields. Plot rows from top to bottom: mean fork length (mm) with standard deviation bars and linear regression line, mean prey organism abundance in salmon stomach contents, mean density of in-field zooplankton (organisms m^-3^), and water temperature (°C) in the middle of a representative field (field 4) for each year 2013-2016 in columns displayed from left to right.

### Overall salmon survival and growth

Survival between years was variable, ranging from 7.4% in 2013 to 61.6% in 2016 and increased over the course of the multi-year experiment (Table 3). A significant increase in survival was observed after 2013 when an undersized culvert in the drainage canal was replaced allowing for much more rapid field draw down in subsequent years.

**Table 3:** Mean percent survival, apparent fork length growth rate and apparent weight growth rate for each year of the experiment.

| Year | Mean survival | Fork length growth rate (mm d<sup>-1</sup>) | Weight growth rate (g d<sup>-1</sup>) |
| --- | --- | --- | --- |
| 2013 | 7.4% | 0.99 | 0.19 |
| 2014 | 44.6% | 0.99 | 0.14 |
| 2015 | 43.5% | 0.81 | 0.12 |
| 2016 | 61.6% | 1.28 | 0.21 |

Juvenile Chinook Salmon apparent growth rates observed in experimental fields were high in all years ranging from 0.81 mm d-1 and 0.12 g d-1 in 2015 to 1.28 mm d-1 and 0.21 g d-1 in 2016 (Table 3). These growth rates were 2-5x higher than previously or concurrently observed in the adjacent Sacramento River [50,51].

## Discussion

Rearing of juvenile Chinook Salmon within winter-flooded rice fields shows strong potential for reconciling agricultural floodplain land use with habitat needs of an imperiled and economically important fish. Winter-flooded rice fields demonstrated high production of naturally occurring fish food (zooplankton) leading to high growth rates of salmon reared in these environments. As with past fish conservation studies in altered environments [52–54], our adaptive research approach enabled us to successfully answer experimental questions despite unpredictable winter hydrologic and temperature regimes in the Central Valley.

In our studies, field substrate did not have a statistically significant effect on the composition or abundance of zooplankton species. Similarly, no significant differences in growth rates of rearing juvenile salmon were detected between post-harvest substrate treatments. Overall, growth across all treatments was extremely fast and much greater than that documented in the Sacramento River channel environments [29]. Accordingly, we do not recommend a specific post-harvest straw management practice. Instead we feel that field preparation should be left to the farmer. We do however, encourage future research that explores other approaches for enhancing in-field habitats to decrease predation risk for rearing fish.

There is currently limited scope for providing avian predation refugia for fish inside of winter-flooded rice fields. We investigated the potential of in-field trenches to provide depth refuge from avian predation, but direct benefits to survival were found to be insignificant in this study. Despite this result, we determined that trenches produced other beneficial by-products. For example, the fields containing perimeter trenches connecting the inlet and outlet structures showed higher rates of volitional emigration of salmon. We speculate that fish used the trenches as migration corridors when emigrating from the fields resulting in a diversification of emigration timing which has been identified as a key component of population stability via the portfolio effect (Carlson and Satterthwaite, 2011) [55]. Additionally, the trenches buffered water temperatures from the daily maximums observed in the middle of the fields and enhanced field drainage with reduced standing water and stranding of fish during field drainage.

In floodplain river ecosystems, fishes often respond strongly to hydrological dynamics of ascending and descending flood conditions [56–58]. Juvenile Chinook Salmon in the Central Valley have evolved physiological and behavioral strategies for the use and egress from winter-flooded floodplain habitats [28,33,59]. Accordingly, the dynamics of draining rice fields containing juvenile Chinook Salmon may matter in terms of providing the appropriate type and quantity of cues to promote the volitional egress of fish from fields with minimal stranding. In our study, extending the drainage period and manipulating inflow conditions had a detrimental effect on survival and the best method was a fast drain where fields were drained in a single day. This was likely the result of increased vulnerability to predation and reduced thermal buffering due to a prolonged period with shallower water depths. Again, these results provide a relatively simple management recommendation for farmers in that a more complicated long drain does not currently appear to be necessary. Rather, simple opening of water control structures combined with volitional passage appears to be the best method. We encourage exploration of other drainage methods, and production of other species in winter-flooded rice fields may require different draining practices.

An initial mortality of approximately 33% was observed in the first week of the 2016 salmon survival over time experiment. The cause of this initial mortality is unknown but was likely a combination of factors, including a stressful transport. Transport is a known stressor on many fishes, including juvenile Chinook Salmon [60,61]. In our study, fish were captured from hatchery raceways, coded wire tagged, allowed to recover for several days and then placed in a fish hauling tank at high densities (up to 25,000 fish m^-3^) and delivered to the fields in early February. Exposing naïve hatchery salmon to a new environment in the flooded agricultural fields may have increased stress as it necessitated behavioral adaptations of prey switching and predator avoidance as well as rapid acclimation to the new physical water quality parameters. After the startlingly high rate of initial mortality, survival stabilized in week 2 and remained high for the remainder of the experiment. Without accurate assessment and accounting of initial post-release mortality there is potential for fishery managers to be chronically overestimating habitat-specific mortality rates determined by recapture of hatchery fish transported and released into natural habitats. We therefore recommend that future research examine effects on initial post-release mortality of transporting, acclimatizing, and releasing hatchery fish into the wild.

There has been a perceived conflict between conservationists and farmers when managing off-season agricultural fields as fish habitats. While farmers have incentives to prepare their fields for a new rice crop as early in the spring as possible, fish conservationists prefer to keep the rice fields wet as late as possible to maximize fish growth and survival before release into the river [32,33]. However, in practice, when weather conditions are good for fish (i.e., wet and cool) in the late winter and early spring, they are generally not conducive to agricultural field preparation. The inverse is also true, when spring conditions are dry and hot and generally suitable for agricultural field preparation, water quality conditions (especially water temperature) are often unsuitable for juvenile Chinook Salmon [62,63]. Thus, given proper timing and coordination within an adaptive management framework, rice farmers and fish conservationists can collaborate to promote threatened fisheries without impacting crop yields [29]. We therefore encourage the continued development of frameworks that work for both fish and farms in floodplain habitats in the Central Valley. In particular, incentive programs for farmers (e.g., through the USDA NRCS program) may be needed to promote these activities to their fullest potential.

Land manager and farmer involvement has generally exceeded expectations in our projects, and we are optimistic about continued stakeholder involvement for several reasons. Given issues with water scarcity in the Central Valley [64,65], the dual-use of rice fields for agriculture and rearing juvenile salmon could establish stronger water security for farmers [66]. Additionally, the reconciling of fish conservation and rice agriculture provides a sustainable method for conducting a service of rice straw decomposition while using natural processes to fuel a productive aquatic food web [27,67]. Salmon reared in the flooded rice fields exhibit among the highest growth rates documented in freshwater in California (Fig 5) [29]. By creating high quality habitat on their fields, farmers can help bolster fish populations by turning small fry into large, healthy smolts rapidly during mid-winter when water temperatures are low, river flows are high and when predators are less active, thus improving salmon survival rates during outmigration [68].

**Fig 5.**
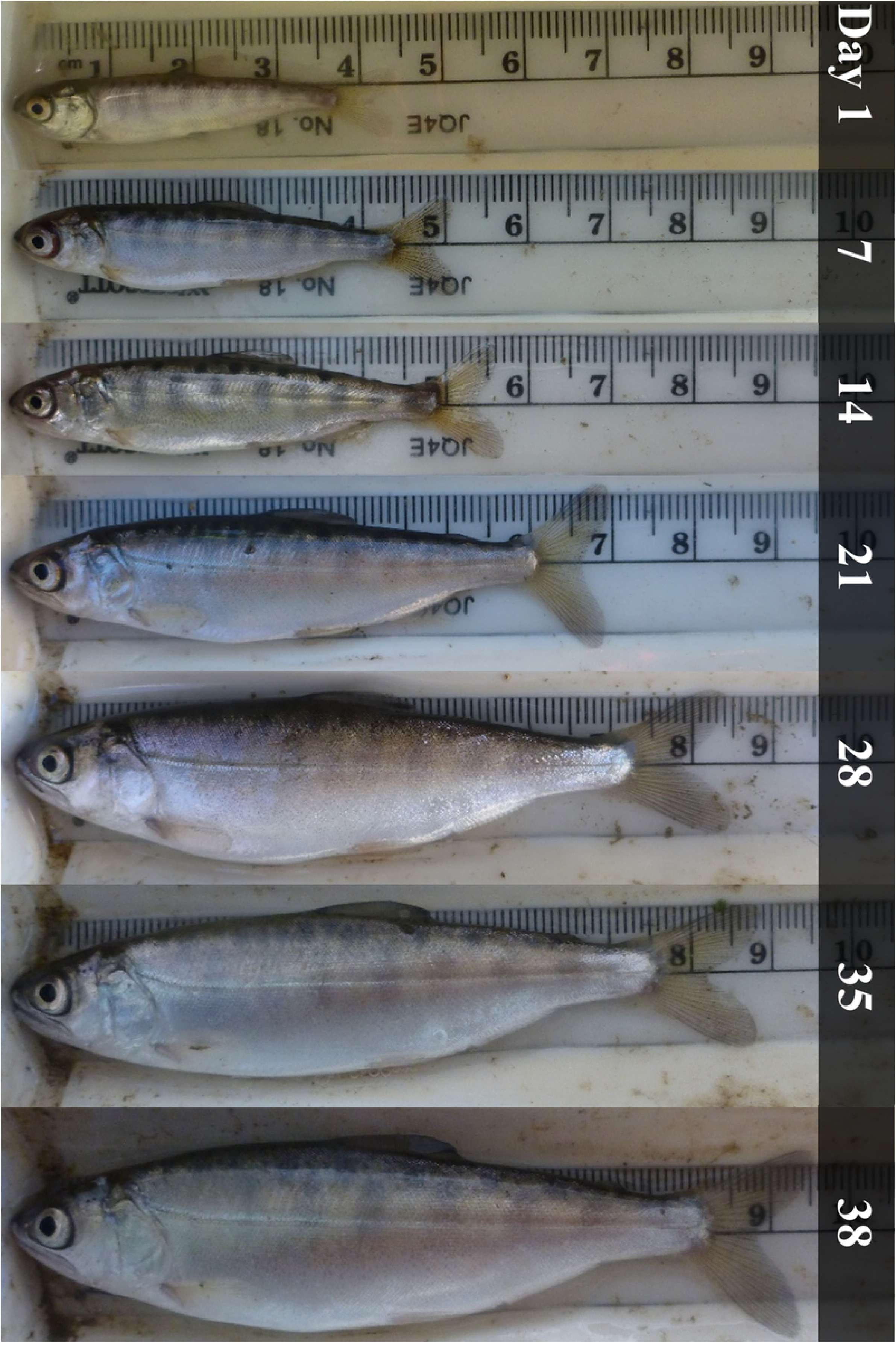
Weekly salmon growth images. Standardized images of a representative juvenile Chinook Salmon sampled weekly during the 2016 experiment.

We support current state and federal programs targeting the modification of existing levee and weir infrastructure to allow fish in river channels to more frequently access managed floodplain habitats during outmigration. We also see the need for additional science to track fish after they leave the fields in order to evaluate the full lifecycle benefits to salmon having reared on inundated agricultural fields (e.g., out-migration survival, smolt-to-adult return rates, and physiological benefits). The exceptional productivity and resulting rapid rates of salmon growth documented on the managed agricultural floodplain highlight the potential value of this habitat for native fish conservation. Managed agricultural floodplains should not be thought of as a replacing the need for preserving the last remnants of natural floodplains, nor should they diminish the conservation need for restoring naturally functioning floodplains where feasible. Managed agricultural floodplains are likely to become an important means for fishery managers to produce ecologically functioning off-channel habits for imperiled native fish especially during times of low water when remaining natural floodplain habitats do not inundate and are therefore inaccessible to salmon populations confined to leveed stream channels.

## Acknowledgements

Salmon used for field experiments were graciously provided by the Feather River State Fish Hatchery. The location for the experiment was provided by Cal Marsh and Farm Ventures with field preparations and maintenance done in collaboration with their farmers. We also thank the California Department of Water Resources and the California Department of Fish and Wildlife for valuable contributions on these studies. We thank a large number of researchers and field technicians who assisted with field experiments through the years, especially Nicholas Corline, Mollie Ogaz, Emma Cox, and Rosa Cox. Funding for the multi-year project was provided through the U.S. Bureau of Reclamation and California Department of Water Resources. PS and ALR were supported through funds from California Trout, Inc., the California Rice Commission, and the USDA NRCS program, Task Order 21 in a master agreement between UC Davis and California Trout Inc. ALR was also supported by the Peter B. Moyle & California Trout Endowment for Coldwater Fish Conservation and the California Agricultural Experimental Station of the University of California Davis, grant number CA-D-WFB-2467-H.

